# Estimating voluntary and involuntary attention in bistable visual perception: A MEG study

**DOI:** 10.1101/2020.02.18.953653

**Authors:** Parth Chholak, Vladimir A. Maksimenko, Alexander E. Hramov, Alexander N. Pisarchik

## Abstract

We introduce a method for measuring human attention when performing a visual task consisting in different interpretations of a bistable image. The Necker cube with flickering faces was presented to nine conditionally healthy volunteers. The pixels intensity in the front and rear cube faces were modulated by a sinusoidal signal with 6.67-Hz (60/9) and 8.57-Hz (60/7) frequencies, respectively. The tags of these frequencies and their second harmonics were clearly identified in the average Fourier spectra of the magnetoencephalographic (MEG) data recorded from the occipital cortex. In the first part of the experiment, the subjects were asked to voluntary control their attention by interpreting the cube orientation as either left- or right-orientated. Accordingly, we observed the dominance of the corresponding spectral component and voluntary attention performance was measured. In the second part of the experiment, the subjects were just asked to observe the cube image without any effort in its interpretation. The alternation of the dominant spectral energies at the second harmonic tag frequencies was treated as changes in the cube orientation. Based on the results of the first experimental stage and using wavelet analysis, we developed a novel method which allowed us to identify currently perceived cube orientations. Finally, we characterized involuntary attention using the dominance time distribution and related it to voluntary attention performance and brain noise. In particular, we have shown that higher attention performance is associated with stronger brain noise.

## Introduction

Wilhelm Wundt was the first who suggested, as early as in 1897, that there exist two forms of attention: voluntary and involuntary [1]. There is already a surplus amount of terms used in the community that overlap with these two forms of attention such as endogenous versus exogenous attention, automatic versus controlled attention, and pull versus push attention [2]. According to Prinzmetal and his colleagues, voluntary and involuntary attention have different functions and are controlled by distinct mechanisms. They supposed that voluntary attention affects perceptual attention and would affect both accuracy and reaction time (RT) experiments, whereas involuntary attention deals with the response selection decision and is manifested only in RT experiments.

To investigate such a distinction, a spatial cuing task developed by Posner and his colleagues [3–6] is useful. In this paradigm, subjects perform a detection or identification task with a peripheral stimulus. The participants are precued to a possible location of the stimulus beforehand; in valid trials, the cue indicates the target location, whereas in the case of invalid trials, the cue indicates a nontarget location. Since the participants are not allowed to move their eyes to the cued location, the observed differences in performance between valid and invalid trials reflect differences in attention which are independent of fixation. Jonides [7, 8] used this paradigm to study the difference between voluntary and involuntary attention by altering the “validity” of the cuing information. If the total number of valid trials for the correct stimulus location is as low as that for a random distribution in which no useful bias for the target location is provided, only involuntary attention would be involved in seeing the peripheral stimulus. On the other hand, in the presence of a high number of valid trials in which correct cuing information for the target location is available, both voluntary and involuntary attentions would be engaged.

Moreover, the effect of involuntary attention on the initial response selection decreases as stimulus onset asynchrony (SOA) increases. The SOA is defined as the time between the onset of cue and target [9, 10]. At the same time, voluntary attention to the biasing cue improves the perceived contrast of attended and unattended stimuli [11–13]. Whether the enhance in the contrast is due to increasing dominance of attended stimulus [14] or decreasing dominance of unattended stimulus [9] is still unclear.

In 2005 Prinzmetal et al. [2] introduced the idea of channel enhancement and channel selection in order to show how the two kinds of attention manifest. Channel enhancement is a process driven by voluntary attention that causes the visual system to gather more information from the attended stimulus than from the unattended stimulus specified by the informative cues. It changes the perceptual representation so that the observers have a clearer view of the stimulus they are attending to [15–17]. The channel enhancement should affect experiments around detection accuracy in the target location which is being attended. Furthermore, it may also improve RT as information is presumably gathered faster in the cued than in the uncued location. Besides, the channel selection deals with the decision making in determining the correct target location or response selection, and would only affect the RT experiments.

There is a general consensus that the Stroop effect alters response selection only, but not perceptual representation [18, 19]. For example, when shown the word BLUE written in red ink and asked the ink color, it would lead to a competition in the response selection that delays the response, but no alteration in the perceived color would be observed. Similarly, involuntary attention will affect RT, but not detection accuracy. Conveniently, several researchers found that involuntary attention to a stimulus only affects the response selection [9, 10, 20].

It should be noted that there is precedence for accuracy and RT studies to produce different effects [21–23]. In particular, Santee & Egeth [23] worked on the redundant target paradigm, in which a target letter can be repeated in the display. They reported that the repeating target speeds up the reaction [24–26], a phenomenon known as a flanker effect, but in turn reduces accuracy [27, 28]. There is evidence that voluntary and involuntary attentions affect SOA differently. Specifically, the SOA in the case of involuntary attention engagement is smaller than for voluntary attention [29, 30].

In this paper, we employ multistable perception as a tool for studying voluntary and involuntary attention. Multistable perception is the phenomenon where the same stimulus can be perceived in two or more different ways. With regards to degrees of freedom, the simplest form of multistable perception is bistable perception, when two different interpretations of the same stimulus are possible. There has been extensive research on this topic over the last two decades and many descriptive models have been proposed [31–36]. The switches between alternative percepts have been proposed to be driven by stochastic processes in the brain [31, 37] due to random neurophysiological activity or neuronal adaptation [34, 35] defined as slow destabilization of currently dominant perception after being active for a prolonged time or due to both noise and adaptation [32, 35, 36]. Each percept competes for dominance over another rival state and the active state tends to suppress alternate perception. Whether the interstate suppression is realized before binocular confluence, such as in the primary visual cortex or the lateral geniculate nucleus [38–40] or after [41, 42] was a matter of numerous debates. The latter mechanism suggests that competition exists between high-level stimulus representations in visual neurons. Behavioral studies [9] support the latter mechanism.

Similarly, the phenomenon of visual attention is based on the competition of one object among a variety of other competing alternatives for enhanced perceptual representation (voluntary attention). This led to the suggestion that bistable perception and attention may be related processes [43, 44]. The previous studies on this topic were performed using an evoked response that consisted of numerous relatively short trials as opposed to a single long trial. On the contrary, in the present work we design experiments to characterize voluntary and involuntary attention using visual responses from relatively long (120-s) trials. Previously, involuntary attention was only found in RT experiments under the evoked response regime. Instead of the evoked response, long entrained visual signals can vary in phase and hence they are unlocked in time with the start of stimulation. Therefore, here we introduce a new term called *visual induced field* (VIF) to distinguish it from the traditional visual evoked field [45].

The first part of our experiment includes a study of controlled (or voluntary) attention on one of the Necker cube orientations, whereas the second part deals with involuntary attention when the subject does not try to control his/her decision about the observed object. We characterize the subjects’ attention ability from the first experiment and then use these results to develop a method based on wavelet energies to measure involuntary attention on actively perceived cube orientation. Lastly, we also characterize involuntary attention using dominance time distributions and make comparisons with voluntary attention and brain noise.

## Materials and methods

### Experimental setup

Magnetoencephalographic (MEG) data were recorded with a whole-head Vectorview MEG system (Elekta AB, Stockholm, Sweden) with 306 channels (102 magnetometers and 204 planar gradiometers). The system was placed inside a magnetically shielded room (Vacuum Schmelze GmbH, Hanau, Germany) at the Laboratory of Cognitive and Computational Neuroscience, Center for Biomedical Technology, Technical University of Madrid, Spain. Fastrak digitizers (Polhemus, Colchester, Vermont) were used to obtain the three-dimensional head shape using approximately 300 points on the scalp of each subject. Additionally, three fiducial points (nasion, left and right preauricular) were acquired for co-registration purposes. A vertical electrooculogram was placed to capture blinks. A single empty room recording of more than two minutes duration was performed on each day of the experiment (Day-1: 4 subjects; Day-2: 5 subjects; Day-3: 3 subjects). Data were sampled at 1000 Hz with an online anti-alias bandpass filter between 0.1 Hz and 330 Hz.

### Participants

Twelve^1^ healthy subjects (aged 17–64 years, 6 males and 6 females) with normal or corrected-to-normal vision participated in the experimental study. All subjects provided written informed consent before the commencement of the experiment. The experimental studies were performed in accordance with the Declaration of Helsinki. Methods were carried out in accordance with approved guidelines. The research was approved by the Ethics Committee of the Technical University of Madrid.

### Visual stimuli

The visual stimulus was a grey Necker cube image on a grey background generated by a personal computer on the computer monitor with a 60-Hz frame rate and projected by a digital light processing projector onto a translucent screen located 150 cm away from the subject. The pixels’ brightness on the left and right cube faces was modulated by a sinusoidal signal with 6.67-Hz (60/9) and 8.57-Hz (60/7) frequencies respectively, as shown in Fig 1. The modulation depth was 100% with respect to the medium greyscale level of the pixels’ brightness (127 in an 8-bit format), i.e., the image brightness varied from black (0) to grey (127). The sinusoidal shape of stimulation and modulation frequencies were chosen in preliminary experiments with other possible flickering frequencies, which are integral fractions of the 60-Hz frame rate (i.e., 60/2, 60/3, 60/4, 60/5, 60/6, 60/7, 60/8, 60/9, 60/10, and 60/12), as frequencies which produce the best tagging brain response [46].

**Fig 1.**
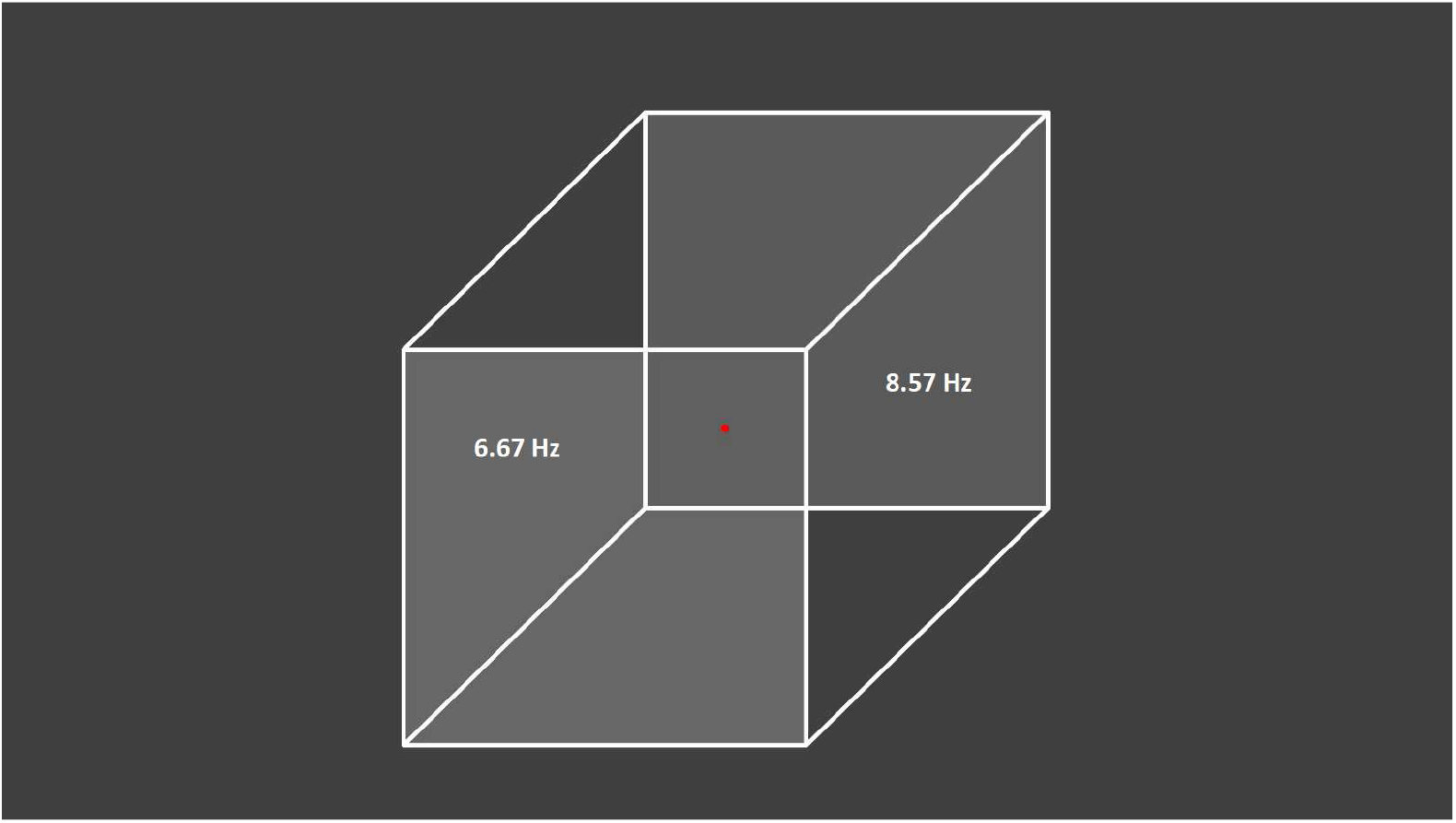
Presented visual stimulus. Necker cube with flickering left and right faces at 6.67 Hz and 8.57 Hz, respectively.

### Experimental procedure

The subjects were sat in a comfortable reclining chair with their legs straight and arms resting on an armrest in front or on their laps. The participants were asked to remove any metallic items above their waist like jewelry, belts, and brassieres, along with their shoes before the experiment. The experiment began with the recording of a two-minute background activity while the subject was focusing on a red dot at the middle of a stationary (non-flickering) cube image. This MEG trial acted as a background reference for further measurements.

The experiment included two stages, voluntary control of the perceived cube orientation and involuntary spontaneous switching between the two cube orientations. During the first stage, after a 30-s rest and an instructional visual message, the flickering Necker cube with two frequencies was presented 24 times on the screen (5-s each with a 5-s interval in-between). For the first 12 trials, 9 out of 12 participants were asked to interpret the cube as left-oriented. After a 30-s rest and an instructional visual message, the participants were requested to interpret the next 12 cubes as right-oriented. For 3 subjects, we reversed the order of voluntary perception by asking them to interpret the first 12 cubes as right-oriented and the next 12 cubes as left-oriented. This concluded the first experimental stage.

When the subject was ready, the second experimental stage started. The same Necker cube stimulus was presented for 120 s. At this stage the subjects were instructed not to fix their attention on a particular cube orientation, but only at the red dot at the centre of the image.

### Visual induced fields (VIF)

The brain is modeled using a mesh of 15004 points representing cortical sources. There are multiple combinations in which these numerous brain sources can produce the observed magnetic activity in the 306 MEG channels. This so-called inverse problem is ill-posed and can only be solved by using additional assumptions about the neuronal system such as minimization of the total energy of the system. The effect of depth-dependent sensitivity and spatial resolution was normalized using the standardized Low-Resolution Electromagnetic Tomography (sLORETA) method.

We used the Brodmann atlas in Brainstorm [47] to find cortical sources associated with visual areas V1 and V2 on the modelled cortical mesh (1227 points). We then averaged the response of these visual sources to obtain VIF for each trial.

### Spectral analysis

Morlet wavelets constructed from a mother wavelet with a 1-Hz central frequency and a 12-s full width at half maximum (FWHM) were utilized to obtain wavelet power time series at second harmonics of the flicker frequencies [46]. The second harmonic frequencies were fine-tuned based on the power spectrum of the VIF signals for each subject.

### Wavelet analysis

The time-frequency analysis is based on the continuous wavelet transform [48, 49]

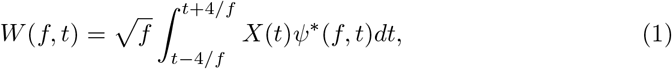

where “*” signifies the complex conjugate and *X*(*t*) is the analyzed MEG signal. The complex-valued Morlet-wavelet is chosen as the mother function

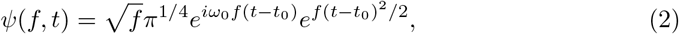

with *ω*_0_ = 2*πf*_0_ being the central frequency of the Morlet wavelets and 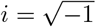. The wavelet powers *W*(*f*_1_, *t*) and *W*(*f*_2_, *t*) given by Eq (1) were evaluated at the tagging frequencies *f*_1_ = 13.33 Hz and *f*_2_ = 17.14 Hz (second harmonics of the flicker frequencies), respectively. Since the frequency response decays with increasing frequency as a 1/*f* rule, the wavelet energy was normalized to the corresponding modulation period (1/*f*_1,2_). Hence, the wavelet time series were multiplied to their defining frequencies to get

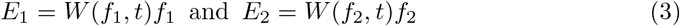

and the difference between the spectral energies at *f*_1_ and *f*_2_ was then calculated as

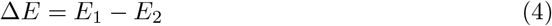

and normalized to its maximum absolute value as

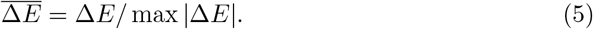

We averaged *E*_1_ and *E*_2_ over time and over all trials separately for the left- (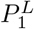 and 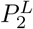) and for the right-oriented cube interpretations (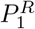 and 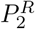). The average spectra are shown in Fig 2.

**Fig 2.**
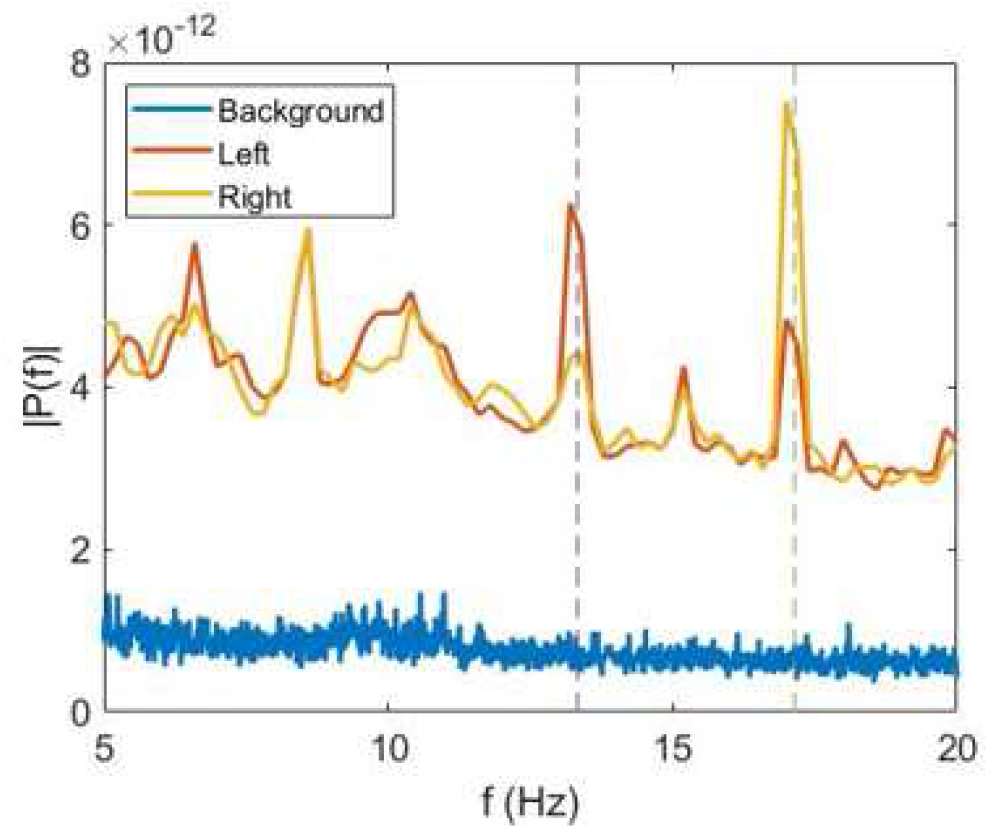
Average power spectra. VIF for all subjects during perception of background, left-oriented cube, and right-oriented cube.

The evolution of the normalized energy difference Eq (5) for typical trials corresponding to the left and right cube orientations for one of the subjects is shown in Fig 3.

**Fig 3.**
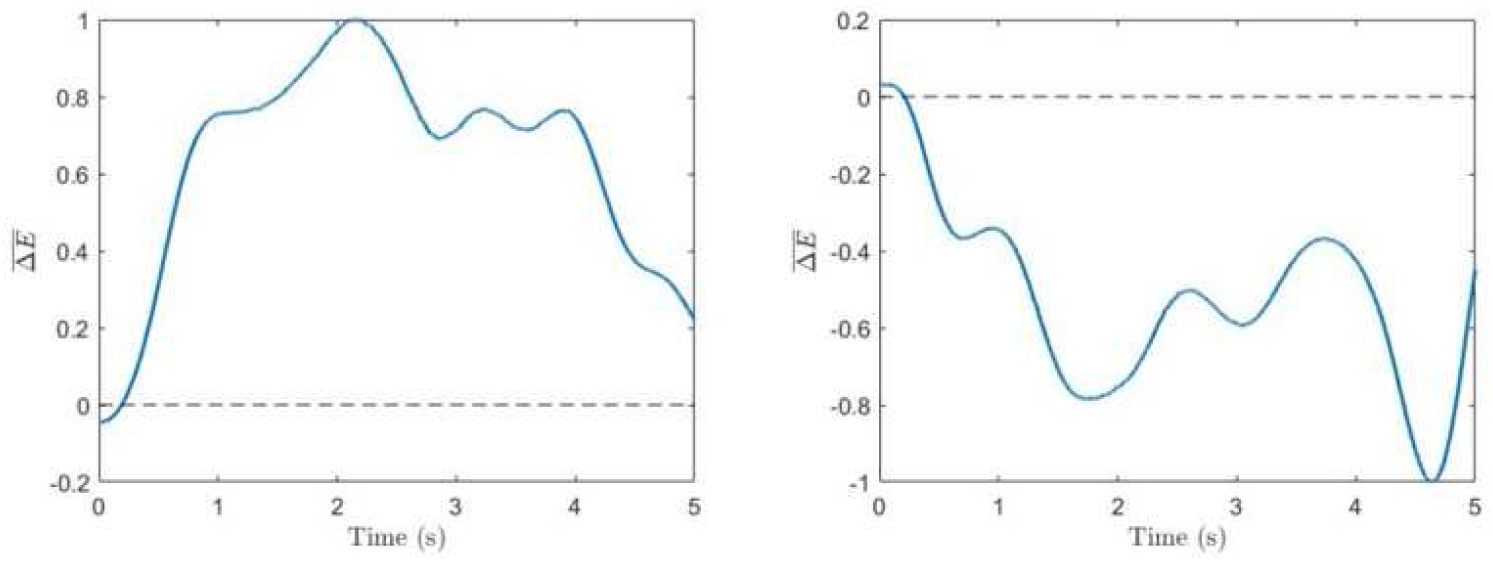
Spectral difference. Time series for single trials corresponding to voluntary left-oriented (left panel) and right-oriented (right panel) cube perception.

The differences between the wavelet energies at *f*_1_ and *f*_2_ corresponding to the left-oriented and right-oriented cube perceptions 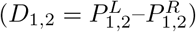 signify the bias in spectral reflection of left orientation in comparison to right orientation such that D1 should be higher and *D*_2_ should be lower. The difference between *D*_1_ and *D*_2_ gave the performance index *μ* as

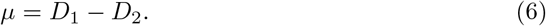

The performance *μ* characterizes the ability of the subject to concentrate voluntary attention. Similar to the voluntary case, wavelet power time series for both frequencies were evaluated from VIF for involuntary perception. However, the time duration of the trials was increased to 120 s.

### Marking perception states

To determine the moment of switches between two different cube orientations, we propose a method based on wavelet power time series. In our approach, 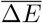 calculated by Eq (5) was screened for significant changes above a threshold equal to its standard deviation *δ*:

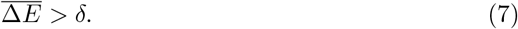

The active state was determined as left-oriented (Switch = 1) if 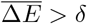 and as right-oriented (Switch = 0) if 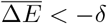. The algorithm is resilient to insignificant perturbations and sticks to the previous state for 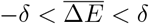. Typical switches in perception between the two cube orientations are illustrated in Fig 3.

**Fig 4.**
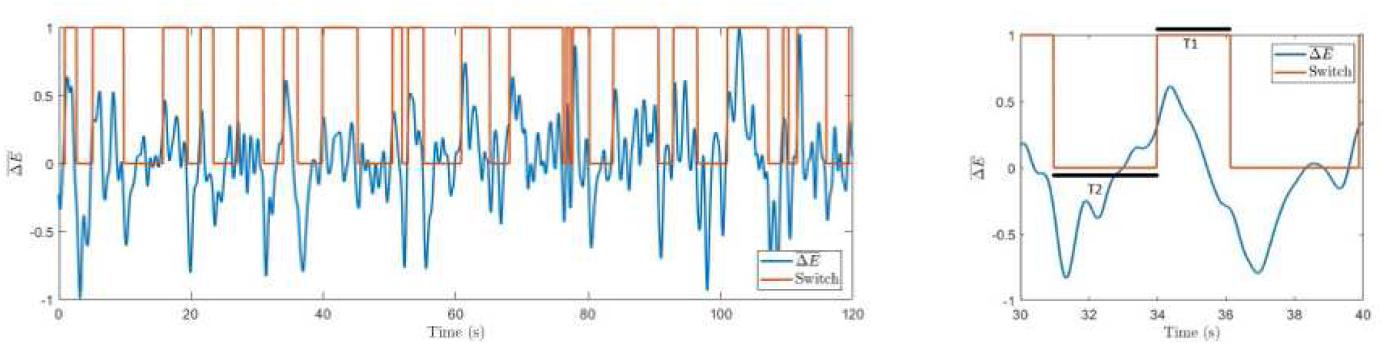
Difference in spectral energies. Time series with marked switches between involuntary left-oriented and right-oriented cube perception. The right panel shows an enlarged part of the left graph with marked resting times T1 and T2.

## Results and discussion

### Experiment-1: Voluntary control of perceived cube orientation

The frequency tagging was successfully used in MEG research for studying perception of ambiguous images [50]. Similarly, in our experiments on voluntary attention, we observed that when the subject endeavored to interpret the cube as left-oriented, the power spectrum at *f*_1_ exhibited higher energy than at *f*_2_, whereas for the right-oriented cube the contribution of *f*_2_ was stronger. This can be seen in Fig 2, where we plot the power spectra averaged over all subjects during the left-oriented cube, right-oriented cube, and stationary cube (or background) without flickering.

Hence, we expect the difference between spectral powers corresponding to the left and the right cube orientations at *f*_1_ (or *D*_1_) to be positive or at least higher than the spectral power difference at *f*_2_ (or *D*_2_) which should either be negative or at least lower than *D*_1_. Furthermore, the difference between *D*_1_ and *D*_2_ would signify the performance in voluntary attention (*μ*) of the subject to tend to both cube orientations, as the cause for perceived contrast between the attended and the unattended stimuli is voluntary attention. Figure 3 shows typical spectral power difference time series for the left and right face frequencies during voluntary attention on left and right cube faces. The spectral difference was largely positive for the left face and negative for the right face as predicted.

For all subjects, we found that *D*_2_ was negative, whereas *D*_1_ was only positive for 9 out of 11 subjects. In Fig 5 we present the voluntary perception performance *μ* calculated by Eq (6) which is always positive, as expected.

**Fig 5.**
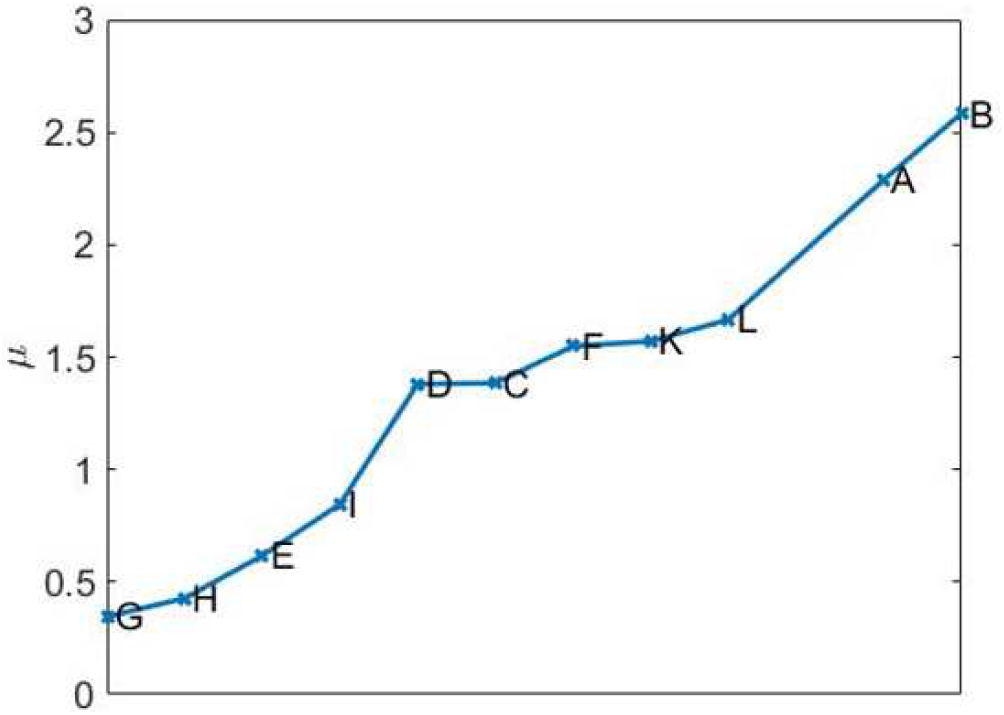
Voluntary attention. Performance of all subjects sorted in ascending order.

### Experiment-2: Involuntary switches between two perceptual states

When the subjects spontaneously switch their attention to either of the cube orientations, the VIF spectral content exhibits narrow peaks at the tagging frequencies *f*_1_ and *f*_2_ and the sum flicker frequencies (*f*_1_ + *f*_2_)/2 (Fig 6. This can be explained by the fact that during perception of either of the cube orientations, the central square at the intersection of both orientations was flickering at the superposition frequency, and subsequently attended during the perception of either orientation.

**Fig 6.**
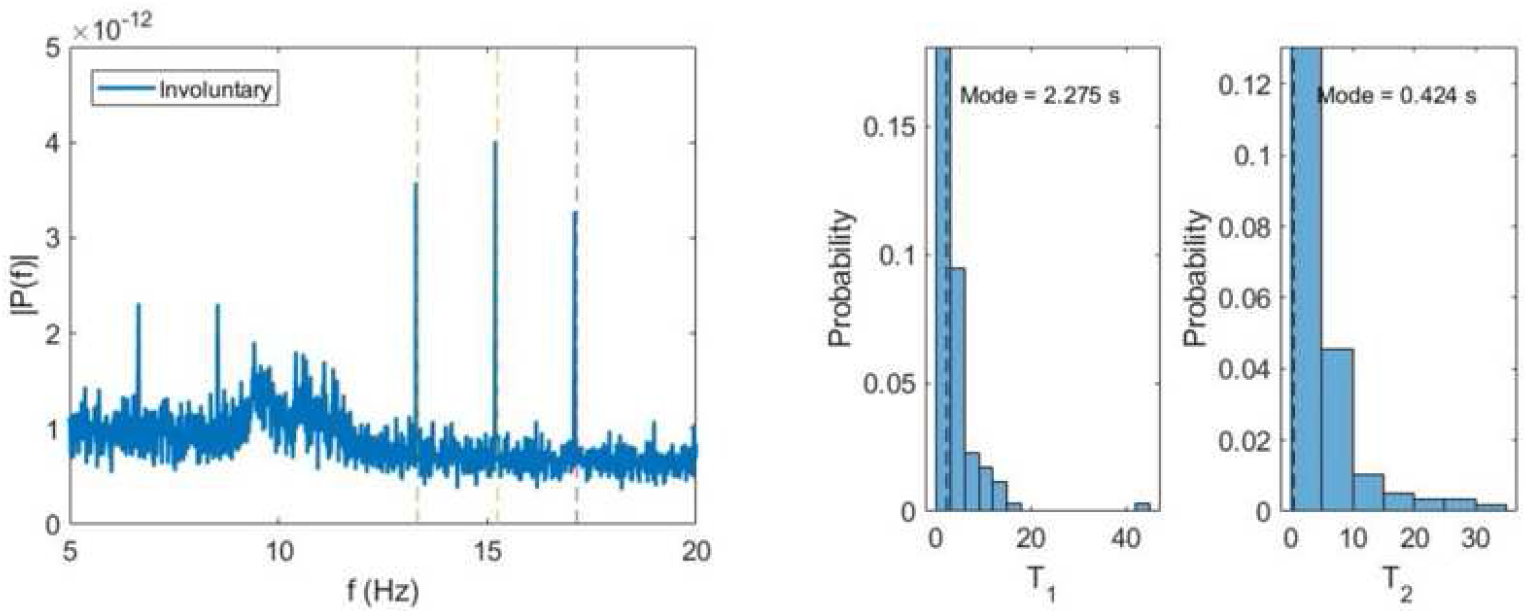
Involuntary attention. (Left) Average power spectrum of VIF for all subjects during involuntary switches between cube orientations. (Middle and right) Probability distributions of dominance times for left (T_1_) and right (T_2_) cube orientations.

The average values of dominance times for both orientations are similar (*T*_*a*1_ = 4.097 ms, *T*_*a*2_ = 5.124 ms), but curiously, the most probable or modal dominance time for the left orientation (*T*_*m*1_ = 2.275 s) is much higher than for the right orientation (*T_m_*_2_ = 0.424 s). This seems to suggest a bias in the perception of the two cube orientations. The same stimulation excites the left orientation more easily and frequently than the right orientation. One of possible reasons for the preference in the left cube orientation may be that in European languages the reading and writing are from left to right. This can explain why in our everyday practice we observe the left-oriented cube more often than the right-oriented cube and hence the perceptual stability of the left-oriented cube is higher than the right-oriented cube one. At the same time, it is also fair to acknowledge that in all the voluntary attention experiments that preceded the involuntary attention experiments, the first 9 subjects were required to first focus on the left orientation invariably. On the other hand, we observed that the change in the order for the last 3 subjects did not affect the modal dominance time values.

In left panel of Fig 7 we plot the average modal dominance time *T*_*m*0_ = (*T*_*m*1_ + *T*_*m*2_)/2 versus voluntary attention performance *μ*. As noted, only 10 out of 12 subjects participated in the second experiment with an additional defaulter. Interestingly, higher attention performance leads to shorter dominance time. This can be explain by the hypothesis that **higher attention requires a larger neuronal network to process information and make a decision**, that in turn increases neural noise since a larger number of synapses and neurons are involved [51]. Finally, stronger brain noise causes more frequent switching between perceptual states or more frequent response selection (involuntary attention) and hence shorter dominance times.

**Fig 7.**
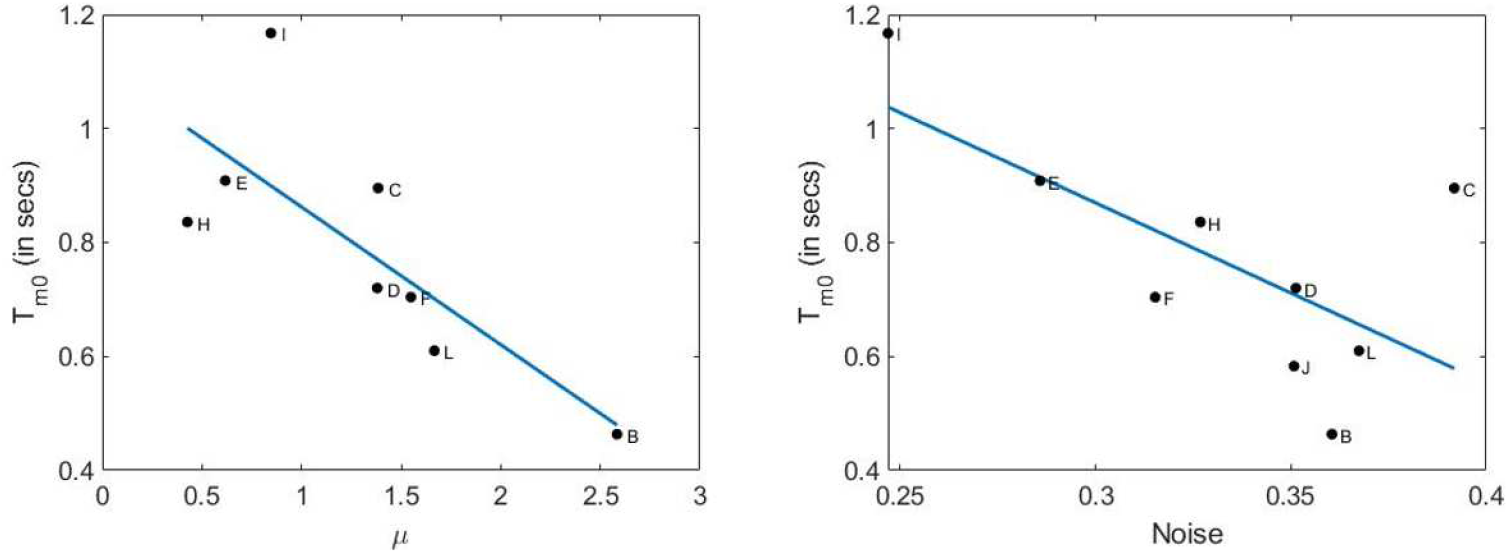
Relation of dominance time with attention performance and brain noise. Left: Dominance time vs attention performance with a linear fit (root mean squared error: 0.168; F-statistics: 5.7; p-value: 0.0484). Right: Dominance time vs brain noise with a linear fit (root mean squared error: 0.147; F-statistics: 8.95; p-value: 0.0242.

To check this hypothesis, we estimated brain noise using the methodology based on phase synchronization [46]. Namely, we measured kurtosis of probability distribution of the phase difference between the second harmonic of the flickering signal (*f*_1_) and visual induced field in the occipital cortex. In the right panel of Fig 7 we plot the average modal dominance time versus brain noise (in units of inverse kurtosis). As expected, these values anticorrelate; this confirms our hypothesis that **higher attention performance is associated with stronger brain noise** because a larger neural network is involved in information processing.

## Conclusion

In this work, we proposed novel approaches for estimating attention performance and classification of bistable perceived states, based on wavelet transformation of neurophysiological brain activity. The developed algorithm for bistable state classification can be useful for designing new noninvasive real-time brain-computer interfaces, due to its fast computation and relative simplicity.

## Acknowledgments

The experimental study was supported by the Spanish Ministry of Economy and Competitiveness under Project SAF2016-80240. The data analysis was supported by the Russian Science Foundation (Grant No. 19-12-00050).

## Footnotes

1 Subject-G and Subject-A did not participate in the involuntary attention experiment due to personal reasons. Additionally, there was a defaulter each in involuntary (Subject-K) and voluntary (Subject-J) attention experiments.

## References

1. Wundt W. Outline of Psychology. Leipzig: Wilhelm Engelmann; 1897.

2. Prinzmetal W, McCool C, Park S. Attention: Reaction time and accuracy reveal different mechanisms. J Exp Psychol. 2005 134(1):73–92.

3. Posner MI, Nissen MJ, Ogden WC. Attended and unattended processing modes: The role of set for spatial location editors’ introduction. In: H. L. Pick and J. E. Saltzman (Eds.), Modes of perceiving and processing information. 1978:137–158.

4. Posner MI. Chronometric explorations of mind. Hillsdale: L Erlbaum; 1978.

5. Posner MI, Snyder CR, Davidson BJ. Attention and the detection of signals. J Exp Psychol. 1980 109(2):160–74.

6. Posner MI. Orienting of attention. Q J Exp Psychol. 1980 32(1):3–25.

7. Jonides J. Towards a model of the mind’s eye’s movement. Can J Psychol. 1980 34(2):103–12.

8. Jonides J. Further toward a model of the Mind’s eye’s movement. Bull Psychon Soc. 1983 21(4):247–50.

9. Hancock S, Andrews TJ. The role of voluntary and involuntary attention in selecting perceptual dominance during binocular rivalry. Perception. 2007 36(2):288–98.

10. Ooi TL, He ZJ. Binocular rivalry and visual awareness: The role of attention. Perception. 1999 28(5):551–74.

11. Carrasco M, Ling S, Read S. Attention alters appearance. Nat Neurosci. 2004 7(3):308–13.

12. Luck SJ. Understanding awareness: One step closer. Nat Neurosci. 2004 7:208–9.

13. Treue S. Perceptual enhancement of contrast by attention. Trends Cogn Sci. 2004 8(10):435–7.

14. Chong SC, Tadin D, Blake R. Endogenous attention prolongs dominance durations in binocular rivalry. J Vis. 2004 5(11):1004–12.

15. Prinzmetal W, Nwachuku I, Bodanski L, Blumenfeld L, Shimizu N. The phenomenology of attention. 2. Brightness and contrast. Consciousness Cogn. 1997 6(2–3):372–412.

16. Prinzmetal W, Wilson A. The effect of attention on phenomenal length. Perception. 1997 26(2):193–205.

17. Prinzmetal W, Amiri H, Allen K, Edwards T. Phenomenology of attention: 1. Color, location, orientation, and spatial frequency. J Exp Psychol. 1998 24(1):261–82.

18. Baldo JV, Shimamura AP, Prinzmetal W. MMapping symbols to response modalities: Interference effects on Stroop-like tasks. Percept Psychophys. 1998 60(3):427–37.

19. Virzi RA, Egeth HE. Toward a translational model of Stroop interference Mem Cogn. 1985 13(4):304–19.

20. Mitchell JF, Stoner GR, Reynolds JH. Object-based attention determines dominance in binocular rivalry. Nature. 2004 429(6990):410–13.

21. Moore CM, Egeth H. How does feature-based attention affect visual processing? J Exp Psychol. 1998 24(4):1296–1310.

22. Mordkoff JT, Egeth HE. Response time and accuracy revisited: Converging support for the interactive race model. J Exp Psychol. 1993 19(5):981–91.

23. Santee JL, Egeth HE. Do reaction time and accuracy measure the same aspects of letter recognition? J Exp Psychol Hum Percept. 1982 8(4):489–501.

24. Eriksen BA, Eriksen CW. Effects of noise letters upon the identification of a target letter in a nonsearch task. Percept Psychophys. 1974 16(1):143–9.

25. Eriksen CW, Eriksen BA. Target redundancy in visual search: Do repetitions of the target within the display impair processing? Percept Psychophys. 1979 26(3):195–205.

26. Eriksen CW, Schultz DW. Information processing in visual search: A continuous flow conception and experimental results. Percept Psychophys. 1979 25(4):249–63.

27. Bjork EL, Murray JT. On the nature of input channels in visual processing. Psychol Rev. 1977 84(5):472–84.

28. Santee J L, Egeth HE. Interference in letter identification: A test of feature-specific inhibition. Percept Psychophys. 1980 27(4):321–30.

29. Posner MI, Cohen Y, Rafal RD. Neural systems control of spatial orienting. Phil Trans R Soc B. 1982 298(1089):187–98.

30. Warner CB, Juola JF, Koshino H. Voluntary allocation versus automatic capture of visual attention. Percept Psychophys. 1990 48(3):243–51.

31. Moreno-Bote R, Rinzel J, Rubin N. Noise-induced alternations in an attractor network model of perceptual bistability. J Neurophysiol. 2007 98(3):1125–39.

32. Shpiro A, Moreno-Bote R, Rubin N, Rinzel J. Balance between noise and adaptation in competition models of perceptual bistability. J Comput Neurosci. 2009 27(1):37–54.

33. Meilikhov EZ, Farzetdinova RM. Bistable perception of ambiguous images: Simple Arrhenius model. Cogn Neurodyn. 2019 13(6):613–21.

34. Dotov DG, Turvey MT, Frank TD. Embodied gestalts: Unstable visual phenomena become stable when they are stimuli for competitive action selection. Atten Percept Psycho. 2019 82(7):2330–42.

35. Huguet G, Rinzel J, Hupe JM. Noise and adaptation in multistable perception: Noise drives when to switch, adaptation determines percept choice. Perception. 2014 14(3):19.

36. Chholak P, Hramov AE, Pisarchik AN. An advanced perception model based on brain noise and adaptation. Nonlin Dyn. 2020 (submitted).

37. Pisarchik AN, Jaimes-Reátegui R, Magallón-García CDA, Castillo-Morales CO. Critical slowing down and noise-induced intermittency in bistable perception. Biol Cybern. 2014 108(4):397–404.

38. Blake R. A neural theory of binocular rivalry. Psychol Rev. 1989 96(1):145–167.

39. Lehky SR, Blake R. Organization of binocular pathways: Modeling and data related to rivalry. Neural Comput. 1991 3(1):44–53.

40. Tong F, Engel SA. Interocular rivalry revealed in the human cortical blind-spot representation. Nature. 2001 411(6834):195–9.

41. Andrews TJ, Purves D. Similarities in normal and binocularly rivalrous viewing. Proc Nat Acad Sci USA. 1997 94(18):9905–8.

42. Logothetis NK, Leopold DA, Sheinberg DL. What is rivalling during binocular rivalry? Nature. 1996 380(6575):621–4.

43. Helmholtz H. Helmholtz’s treatise on physiological optics. New York: Dover Publications; 1962.

44. Leopold DA, Logothetis NK. Multistable phenomena: Changing views in perception. Trends Cogn Sci. 1999 3(7):254–64.

45. Regan D. Human brain electrophysiology: Evoked potentials and evoked magnetic fields in science and medicine. New York: Elsevier; 1989.

46. Pisarchik AN, Chholak P, Hramov AE. Brain noise estimation from MEG response to flickering visual stimulation. Chaos, Soliton Fract X. 2019 1:100005.

47. Tadel FF, Baillet S, Mosher JC, Pantazis D, Leahy RM. Brainstorm: A user-friendly application for MEG/EEG analysis Computl Intel Neurosc. 2011 2011:879716.

48. Hramov AE, Koronovskii AA, Makarov VA, Pavlov AN, Sitnikova E. Wavelets in Neuroscience. New York: Springer Heidelberg; 2015.

49. Pavlov AN, Hramov AE, Koronovskii AA, Sitnikova EY, Makarov VA, Ovchinnikov AA. Wavelet analysis in neurodynamics. Phys-Uspekhi. 2012 55(9):845–75.

50. Parkkonen L, Andersson J, Haämälälinen M, Hari R. Early visual brain areas reflect the percept of an ambiguous scene. Proc Nac Acad Sci. 2008 105:20500–4.

51. Pisarchik AN, Maksimenko VA, Andreev AV, Makarov VV, Zhuravlev MO, Frolov NS, Runnova AE, Hramov AE. Coherent resonance in the distributed cortical network during sensory information. Sci Rep. 2019 9:18325.

